# Alternative phage-host interaction in *Lactococcus lactis*: the carrier state drives rapid evolution of phages

**DOI:** 10.1101/2020.04.27.063958

**Authors:** Barbara Marcelli, Anne de Jong, Thomas Janzen, Jan Kok, Oscar P. Kuipers

## Abstract

*Lactococcus lactis* is a lactic acid bacterium widely used as starter culture for the manufacture of fermented milk products like quark, buttermilk and cheese. Bacteriophage infection of starter cultures is one of the biggest causes of fermentation failure and, therefore, lactococcal phages have received great attention from the scientific community in the past decades. In this work we present evidence for the establishment of a carrier state life cycle (CSLC) by a bacteriophage belonging to the c2 species, in the model laboratory strain *L. lactis* MG1363. Our results show that infection of *L. lactis* MG1363 with a second, dissimilar, c2 bacteriophage can induce the CSLC phage to enter an active lytic life cycle. The viral progeny obtained after this infection is a mixed population of phages with differences in their genome sequences and host ranges, indicative of an extremely rapid evolution process. We discuss the possible implications of this phage-host interaction, both with respect to bacteriophage evolution and phage adaptation to different hosts.

**IMPORTANCE:** Our results broaden the current know-how on the yet poorly investigated phage-host interaction mechanism of CSLC, propose a new bacteriophage evolution mechanism, and demonstrate that the outcome of phage infections is possibly more intricate than presently acknowledged.

## Introduction

Bacteriophages, or phages, are viruses that can only infect bacterial cells. In order to assemble and release a new viral progeny, they hijack the replication, transcription and translation machinery of their host, utilizing it as a factory to synthesize several copies of the viral genome and viral proteins. Bacteriophages are the most abundant biological entities in the biosphere and can be found in any biological niche, where they co-evolve with their bacterial host in a continuous struggle for survival (1–3).

To proliferate inside their sensitive host population, bacteriophages can employ different life cycles; the two most common ones are the lytic and the lysogenic cycle (4). In the first case, typical of strictly lytic bacteriophages, the virus replicates inside the bacterium resulting in most of cases in the burst of the host cell and the release of the phage progeny (5). The lysogenic cycle is typical of temperate phages. In this case the phage is able to choose between active lytic propagation and a lysogenic state in which the viral genome integrates via site-specific recombination into the host chromosome and replicates together with it for many generations (6). Bacteriophages in this state take the name of prophages. An external stress factor such as UV irradiation or antibiotic treatment can induce excision of the prophage from the host chromosome and initiation of its lytic replication cycle (6). However, two less investigated types of bacteriophage replication exist, namely pseudolysogeny and carrier state life cycle (CSLC), two terms that have been used ambiguously in the past (4, 6). Pseudolysogeny describes a situation in which, in case of host starvation, a temperate bacteriophage does not commit to either the lysogenic or the lytic cycle but is, instead, present as an episome inside the host cell. This allows the phage to be replicated, albeit at lower frequency than in a classic lytic cycle, and guarantees phage survival during harsh conditions (5). Normal bacteriophage replication is restored once host starvation is resolved (7). CSLC is a state in which a bacterial population and its lytic bacteriophage population are in a dynamic equilibrium. Even though the majority of the bacteria is resistant to the phage, a small sensitive subpopulation of bacteria exists that is constantly infected and produces a small number of intact phage particles (5, 8). As a result, bacteriophages are constantly released into the environment allowing both phage and host to thrive. CSLC has been described to also occur under optimal host growth conditions (5). Different beneficial effects of CSLC have been reported for different bacterial species, such as better adaptability to non-favorable conditions and bacteriophage superinfection immunity (9–11).

*Lactococcus lactis* is a lactic acid bacterium widely used as starter culture for the production of different fermented foods, especially a large variety of cheeses (12). Bacteriophages infecting *L. lactis* have been intensely investigated in the last four decades as they represent one of the biggest causes of dairy fermentation failure (13). Ten different bacteriophage species are known to infect *L. lactis*, and members of three of them, namely c2, 936 and P335, are the most commonly isolated phages in dairy environments (14–16). The occurrence of CSLC phages in mesophilic complex starter cultures of *L. lactis* and *Leuconostoc mesenteroides,* utilized for the production of fermented milk products, has been previously described and was postulated to play an important role in maintaining the compositional stability of these mixed bacterial communities (17). However, the evolutionary implication of CSLC for bacteriophages has been poorly investigated.

Here we present a case of a CSLC between a lytic phage belonging to lactococcal phage species c2 and the *L. lactis* laboratory strain MG1363. Our data show that, after infection of *L. lactis* MG1363 with a second but different c2 bacteriophage, the CSLC phage is able to enter an active lytic replication cycle and to produce new progeny resulting in visible lysis plaques. As a result, one single infection event generates a mixed population of phages, the members of which differ in their genome sequence, host range, and titration suggesting an extremely fast evolution process.

## Materials and Methods

### Bacterial strains, phages and culture conditions

The bacterial strains and bacteriophages used in this study are listed in Tables 1 and Table 2, respectively. All *L. lactis* strains were grown at 30 °C in M17 liquid medium (BD - Becton, Dickinson and Company, Franklin Lakes, NJ) supplemented with 0.5 % glucose (GM17).

**Table 1.** List of strains used in this study.

| ***L. lactis* strain** | **Subspecies** | **Description** | **Reference** |
| --- | --- | --- | --- |
| MG1363 | *cremoris* | Laboratory model strain. Plasmid-free derivative of the dairy isolate NCDO712 | (30),<br>Genbank:WJVF00000000 |
| AM1 | *cremoris* | Dairy starter strain | (31) |
| SMQ 86 | *cremoris* | Dairy starter strain | (32) |
| SMQ 384 | *cremoris* | Dairy starter strain | (33) |
| SMQ 358 | *cremoris* | Dairy starter strain | (33) |
| SMQ 450 | *cremoris* | Dairy starter strain | (14) |
| SMQ 562 | *cremoris* | Dairy starter strain | (14) |
| ML8 | *cremoris* | Dairy starter strain | (34) |
| 3107 | *cremoris* | Dairy starter strain | (35) |
| 158 | *cremoris* | Dairy starter strain | (36) |
| UC509.9 | *cremoris* | Dairy starter strain | (37) |
| SK11 | *cremoris* | Dairy starter strain | (38) |
| 184 | *lactis* | Dairy starter strain | (37) |
| UL8 | *lactis* | Dairy starter strain | (37) |
| IL1403 | *lactis* | Laboratory model strain. Plasmid-free derivative of the dairy isolate CNRZ157 | (37) |
| C10 | *lactis* | Dairy starter strain | (37) |
| 229 | *lactis* | Dairy starter strain | (37) |
| Bu2 - 60 | *lactis* | Dairy starter strain | (39) |
| CH_LC31 | *lactis* | Dairy starter strain | This work |

**Table 2.**
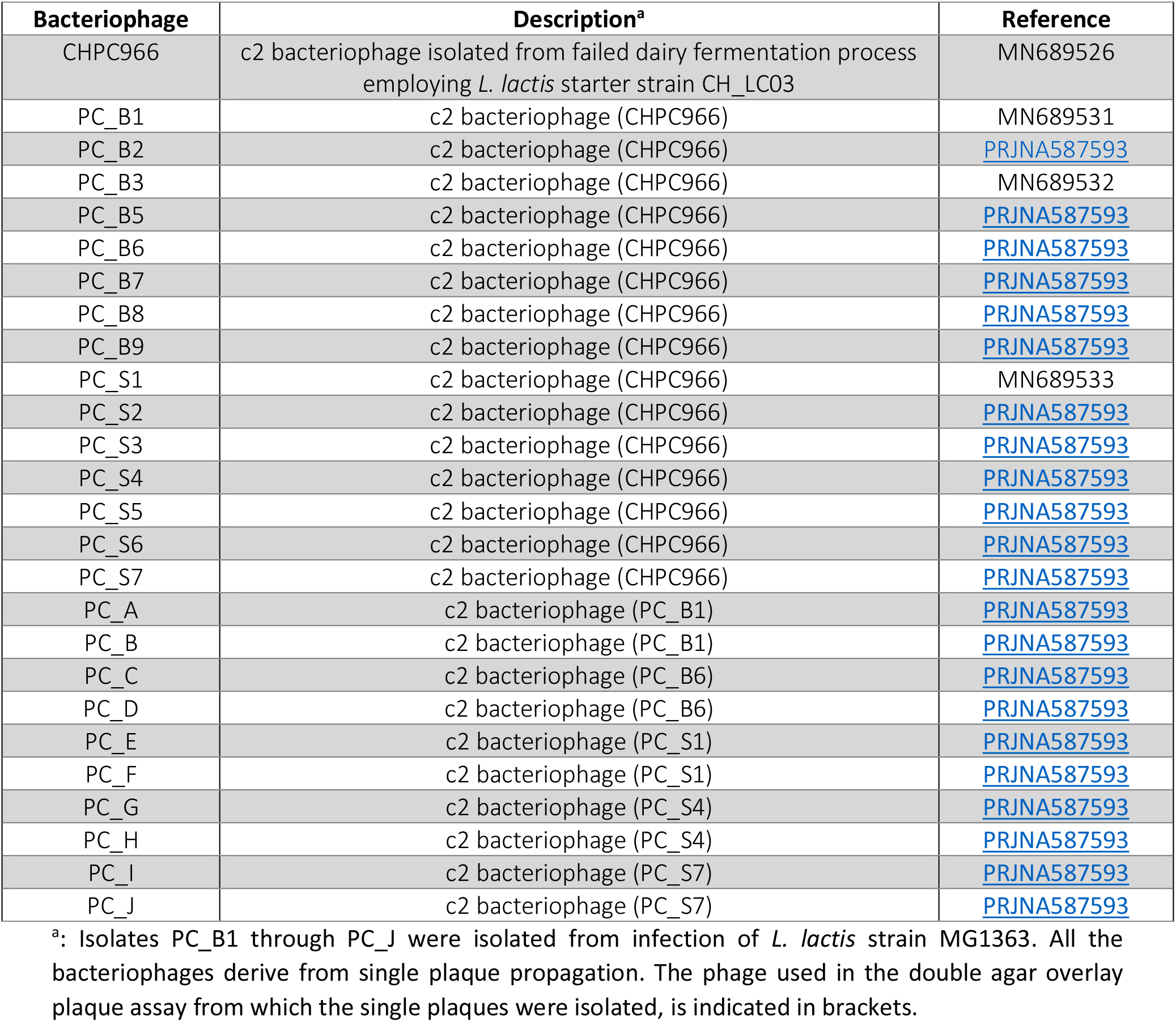
List of bacteriophages used in this study.

| Bacteriophage | Description<sup>a</sup> | Reference |
| --- | --- | --- |
| CHPC966 | c2 bacteriophage isolated from failed dairy fermentation process employing *L. lactis* starter strain CH_LC03 | MN689526 |
| PC_B1 | c2 bacteriophage (CHPC966) | MN689531 |
| PC_B2 | c2 bacteriophage (CHPC966) | PRJNA587593 |
| PC_B3 | c2 bacteriophage (CHPC966) | MN689532 |
| PC_B5 | c2 bacteriophage (CHPC966) | PRJNA587593 |
| PC_B6 | c2 bacteriophage (CHPC966) | PRJNA587593 |
| PC_B7 | c2 bacteriophage (CHPC966) | PRJNA587593 |
| PC_B8 | c2 bacteriophage (CHPC966) | PRJNA587593 |
| PC_B9 | c2 bacteriophage (CHPC966) | PRJNA587593 |
| PC_S1 | c2 bacteriophage (CHPC966) | MN689533 |
| PC_S2 | c2 bacteriophage (CHPC966) | PRJNA587593 |
| PC_S3 | c2 bacteriophage (CHPC966) | PRJNA587593 |
| PC_S4 | c2 bacteriophage (CHPC966) | PRJNA587593 |
| PC_S5 | c2 bacteriophage (CHPC966) | PRJNA587593 |
| PC_S6 | c2 bacteriophage (CHPC966) | PRJNA587593 |
| PC_S7 | c2 bacteriophage (CHPC966) | PRJNA587593 |
| PC_A | c2 bacteriophage (PC_B1) | PRJNA587593 |
| PC_B | c2 bacteriophage (PC_B1) | PRJNA587593 |
| PC_C | c2 bacteriophage (PC_B6) | PRJNA587593 |
| PC_D | c2 bacteriophage (PC_B6) | PRJNA587593 |
| PC_E | c2 bacteriophage (PC_S1) | PRJNA587593 |
| PC_F | c2 bacteriophage (PC_S1) | PRJNA587593 |
| PC_G | c2 bacteriophage (PC_S4) | PRJNA587593 |
| PC_H | c2 bacteriophage (PC_S4) | PRJNA587593 |
| PC_I | c2 bacteriophage (PC_S7) | PRJNA587593 |
| PC_J | c2 bacteriophage (PC_S7) | PRJNA587593 |

<sup>a</sup>: Isolates PC\_B1 through PC\_J were isolated from infection of *L. lactis* strain MG1363. All the bacteriophages derive from single plaque propagation. The phage used in the double agar overlay plaque assay from which the single plaques were isolated, is indicated in brackets.

Bacteriophages were propagated by infecting a 10 ml culture of the indicator strain in its early exponential growth phase (OD_600_ = 0.3 – 0.5) in GM17 containing 10 mM CaCl_2_ with a single plaque. The sample was incubated at 30 °C until visible cell lysis had occurred and subsequently centrifuged at 3,500 x g for 10 min in an Eppendorf tabletop centrifuge 5804R (Eppendorf, Hamburg, Germany) to eliminate cell debris. The supernatant was filter-sterilized using a 0.45 μm filter (GE Healthcare Bio-Sciences, Pittsburg, PA) and stored at 4 °C until use.

Phage titration and plaque morphology assessment were carried out via double agar overlay plaque assay as previously described (4) with the following modifications: bottom and top agar layers contained 1% and 0,4 % agar, respectively. CaCl_2_ was added to the media at the final concentration of 10 mM. Glycine was added at the final concentration of 0.5 % (wt/vol) to facilitate plaque visualization as previously reported (18). Bacteriophage lysates were diluted in TBT buffer (100 mM NaCl, 50 mM Tris-HCl, 10 mM MgCl_2_, [pH7]). 200 μl of an overnight sample of the bacterial strain and 10 μl of properly diluted phage lysate was mixed with 4 ml of top agar, the mix was poured on a solidified bottom agar plate. The plates were incubated overnight at 30 ̊C and subsequently examined for the presence of phage-derived plaques.

Species assignment of bacteriophages was done using multiplex PCR as previously described (19, 20).

### Bacteriophage host range assay

Bacteriophage isolates were tested against an array of *L. lactis* industrial and laboratory strains via spot tests via a double agar overlay assay as previously described. 100-fold diluted samples of the phage lysates (in TBT buffer) obtained as described above were spotted on a solidified top agar layer containing 200 μl of an overnight sample of the bacterial strain. The plates were incubated overnight at 30° C and then checked for the presence of lysis halos in the spot area. In the case of the appearance of spots of clearance, the strain under study was considered as possibly sensitive to the phage. All tests were conducted in triplicate.

### Total bacteriophage and bacterial DNA isolation

Phage DNA was isolated from 5 ml of phage lysate obtained as described above. The lysate was mixed with 10% w/v polyethylene glycol MW 800 and 0.5 M NaCl, and incubated for 16 h at 4°C to allow the phage particles to precipitate. The sample was subsequently centrifuged for 1 h at 11,000 x g at 4°C in an Eppendorf tabletop centrifuge 5810R (Eppendorf). The phage pellet was resuspended in 400 μl DNAseI buffer (10 mM Tris-HCl, pH 7.6, 2.5 mM MgCl_2_, 0.5 mM CaCl_2_). Residual host DNA and RNA were degraded by incubation at 37°C for at least 30 min with 1 μg/ml each of DNAseI and RNAseI (Merk KGaA, Darmstadt, Germany). Ethylendiaminetetraacetic acid (EDTA) was then added at a final concentration of 5 mM and the sample was incubated at 65°C for 15 min to inactivate the enzymes. Phage capsids were degraded by incubating the mixture for 20 min at 56°C with proteinase K (Merk KGaA) at a final concentration of 2 μg/ml. Sodium dodecyl sulfate (SDS) was added to 2.5% v/v and incubation was continued at 65°C for 10 min. Phage DNA was purified by two consecutive phenol/chloroform extractions: the sample (approx. 500 μl) was mixed with an equal volume of a mixture of phenol:chloroform:isoamyl alcohol (25:24:1) in a 2-ml 5-Prime phase lock gel-light tube (Quanta BioScience, Beverly, MA, USA) and centrifuged at 13,000 x g for 8 min in an Eppendorf tabletop centrifuge 5418. The upper DNA-containing aqueous phase was mixed by inversion with 0.1 volume of 3 M sodium acetate (pH 4.8) and 2.5 volumes of 80% ice-cold ethanol and incubated at −20°C for at least 1 h to allow DNA precipitation. DNA was concentrated by centrifugation at 13,000 x g for 15 min in an Eppendorf tabletop centrifuge 5418 and washed twice with 1 ml of 80% ice-cold ethanol. The DNA pellet was air dried for at least 1 h before resuspending in 50 μl TE buffer (10 mM Tris-HCl, pH8, 1 mM EDTA). DNA samples were stored at 4°C until use.

Total bacterial DNA was isolated from 4 ml of an overnight-grown culture. The sample was centrifuged at 14,000 x g for 1 minute on an Eppendorf tabletop centrifuge 5810R (Eppendorf) and resuspended in 300 μl of Birnboim solution A (20 % sucrose, 10 mM Tris HCl pH 8, 10 mM EDTA, 50 mM NaCl) containing 5 mg/ml of lysozyme and 10 mg/ml of RNase. The sample was incubated at 37°C for 30 min to initiate bacterial cell wall hydrolysis and subsequently at 55°C for 10 min to degrade RNA. Bacterial cells were lysed by adding 500 μl of Nuclei lysis solution (Component of Wizard^®^, Wizard^®^ SV and Wizard^®^ SV 96 Genomic DNA Purification Systems, Promega, Madison WI, USA) and incubating at 80°C for 5 min. Subsequently 200 μl of Protein lysis solution (Component of Wizard^®^, Wizard^®^ SV and Wizard^®^ SV 96 Genomic DNA Purification Systems available as a standalone item, Promega, Madison WI, USA) was added and the sample was incubated on ice for 10 min. The suspension was then centrifuged for 10 min at 14,000 x g in an Eppendorf tabletop centrifuge 5810R and the supernatant containing the DNA was transferred to a clean 2 ml Eppendorf tube containing 250 μl of demineralized water and 250 μl of phenol. The sample was mixed by inversion and incubated at room temperature for 10 min before adding 250 μl of chloroform. The water/phenol/chloroform mixture was transferred to a 2-ml 5Prime phase lock gel-heavy tube (Quanta BioScience, USA) and centrifuged for 10 min at 15,000 x g in an Eppendorf tabletop centrifuge 5810R. The upper water-phase obtained from centrifugation was then transferred to a new 2-ml 5Prime phase lock gel-heavy tube already containing 500 μl of chloroform and mixed by inversion. The sample was centrifuged for 10 min at 15,000 x g in an Eppendorf tabletop centrifuge 5810R. The water-phase was transferred to a clean 2ml Eppendorf tube and DNA was precipitated by adding 100 μl of NaAc, pH 5.2 and 700 μl of isopropanol. The sample was mixed by inversion until precipitated DNA was visible and then centrifuged at 15,000 x g for 5 min in an Eppendorf tabletop centrifuge 5810R. The supernatant was discarded and the DNA pellet was washed twice with 1 ml of 70 % of ice-old ethanol. The DNA pellet was then air-dried for 15 min at room temperature and finally dissolved in 100 ul of TE buffer. DNA samples were stored at 4 °C until use.

### Bioinformatics analyses

The nucleotide sequences of the genome of *L. lactis* MG1363 and of the genomes of the bacteriophages studied here were determined using the Illumina MiSeq platform with 2 x 150-bp paired-end sequencing (Illumina, San Diego, CA, USA). Nucleotide sequences were trimmed using Trimmomatic (21) and assembled using A5-myseq (22). The assembled contigs were annotated using the RASTtk server (23). Alignment of the raw sequences contained in the fastq files of *L. lactis* MG1363 total genome sequencing against bacteriophage B3 genome, was performed using bowtie (24). Alignment of the separate sequencing scaffolds against the sequences of phage CHPC966 and those of the representative CSLC phage was performed using BLAST (25) and subsequently visualized using SnapGene^®^ software (from GSL Biotech; available at snapgene.com)

### Accession numbers

The complete genome sequences of bacteriophages CHPC966, PC_B1, PC_B3, and PC_S1 are available at GenBank under the accession numbers: MN689526, MN689531, MN689532, and MN689533, respectively. The whole genome shotgun sequences of all the other bacteriophages studied here are available at the GenBank under the BioProject accession number: <u>PRJNA587593</u>

The whole genome shotgun sequences of *L. lactis* MG1363 strain used here, are available at the GenBank under the accession number: WJVF00000000.

## Results

### Infection of *L. lactis* MG1363 with phage CHPC966 produces a mixed population of bacteriophages

*L. lactis* bacteriophage CHPC966 was isolated in 2002 in a cheese factory based in the USA, from a failed fermentation whey sample (Table 2). To analyze its characteristics, a pure phage lysate was obtained by single plaque propagation on the industrial *L. lactis* strain CH_LC31 employed in the failed fermentation. First, its species identity was determined via multiplex PCR approaches (19, 20), revealing that the phage belongs to the lactococcal phage species c2 (data not shown). Subsequently, its host range was assessed against a panel of lactococcal dairy-derivative and laboratory strains. As shown in Table 3, phage CHPC966 was able to propagate only on strains CH_LC31 and MG1363. Interestingly, in a double agar overlay assay using these two strains, phage CHPC966 produced plaques with different morphologies. On strain CH_LC31, it formed only plaques with a clear lysis center of 1 mm diameter, while on strain MG1363 two plaque types were observed: those with a clear lysis center of 3 mm diameter and a turbid lysis halo, and clear plaques with a 1-mm diameter (Figure 1). On strain MG1363, while the number of plaques with a 1 mm morphology was in agreement with the corresponding lysate dilution used in the assay, the number of plaques of 3 mm remained practically unchanged with ≤ 50 plaques per plate regardless of the phage lysate dilution used (data not shown). The same phenomenon was observed when using stocks of *L. lactis* MG1363 from different sources. This result, combined with the heterogeneous plaque morphologies on infecting *L. lactis* MG1363, suggested that the phage population could be composed of different bacteriophages. To examine this hypothesis, a total of 16 plaques were isolated from four independent propagations of phage CHPC966 on *L. lactis* MG1363. Of these, 9 plaques showed the 3 mm diameter morphology, while the remaining 7 presented a 1 mm diameter morphology. The phages of the 16 plaques were propagated separately on liquid cultures of *L. lactis* MG1363 and the resulting lysates were used to assess the host range of all the isolates. The lysates were named PC_B1 through PC_B9 for those deriving from 3 mm (Big) plaques, and PC_S1 through PC_S7 for those originating from 1 mm (Small) plaques. For reasons of readability, we will refer to these isolates as B1 through B9, and S1 through S7.

**Figure 1.**
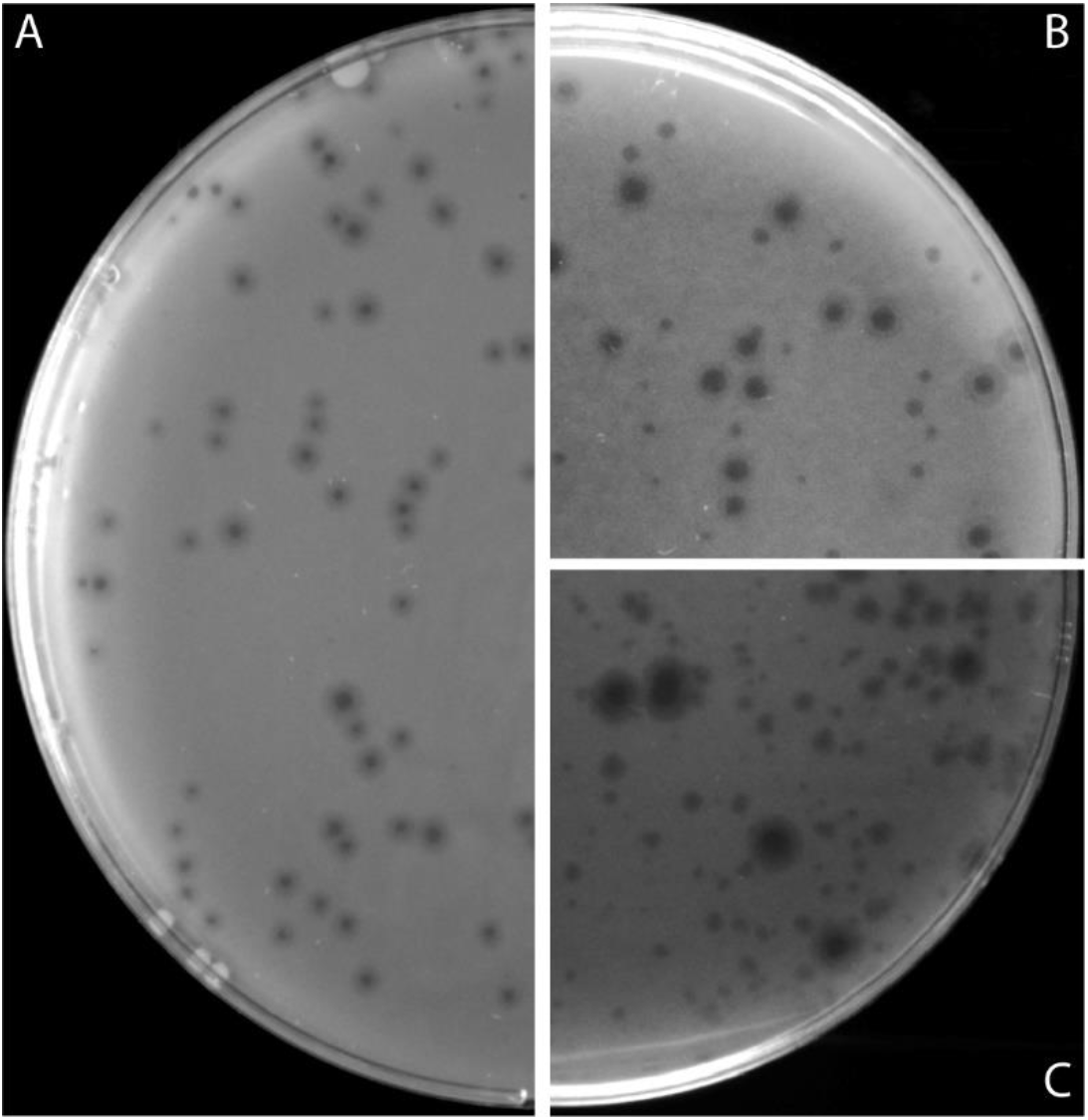
Phage plaque morphology. Panel A: Morphology of plaques obtained after infection of *L. lactis* CH_LC03 with phage CHPC966. Panels B and C: Morphology of plaques obtained after infection of *L. lactis* MG1363 with phage CHPC966.

**Table 3.** Host range of the bacteriophages used in this study.

| Bacteriophage |  | *Lactococcus lactis* strain<sup>c</sup> |  |  |  |  |  |  |  |  |  |  |  |  |  |  |  |  |  |  |
| --- | --- | --- | --- | --- | --- | --- | --- | --- | --- | --- | --- | --- | --- | --- | --- | --- | --- | --- | --- | --- |
| First infection<sup>a</sup> | Second infection<sup>b</sup> | CH_LC03 | MG1363 | 3107 | IL1403 | C10 | ML8 | Bu260 | SMQ384 | AM1 | SK11 | UC509.9 | SMQ450 | SMQ562 | SMQ86 | SMQ385 | 158 | UL8 | 184 | 229 |
| CHPC966 |  | + | + |  |  |  |  |  |  |  |  |  |  |  |  |  |  |  |  |  |
| PC_B1 |  |  | + | + | + | + |  |  | + |  |  |  |  |  |  |  |  |  |  |  |
|  | PC_A |  | + | + | + | + |  |  | + | + | + |  |  |  |  |  |  |  |  |  |
|  | PC_B |  | + | + | + | + |  |  | + | + | + |  |  |  |  |  |  |  |  |  |
| PC_B2 |  |  | + | + | + | + | + | + | + |  |  |  |  |  |  |  |  |  |  |  |
| PC_B3 |  |  | + | + | + | + | + | + | + |  |  |  |  |  |  |  |  |  |  |  |
| PC_B5 |  |  | + | + | + | + | + | + | + |  | + |  |  |  |  |  |  |  |  |  |
| PC_B6 |  |  | + | + | + | + | + | + | + |  | + |  |  |  |  |  |  |  |  |  |
|  | PC_C |  | + | + | + | + | + | + |  | + | + |  |  |  |  |  |  |  |  |  |
|  | PC_D |  | + | + | + | + | + | + |  | + | + |  |  |  |  |  |  |  |  |  |
| PC_B7 |  |  | + | + | + | + | + | + | + |  | + |  |  |  |  |  |  |  |  |  |
| PC_B8 |  |  | + | + | + | + | + | + | + |  | + |  |  |  |  |  |  |  |  |  |
| PC_B9 |  |  | + | + | + | + | + | + | + |  | + |  |  |  |  |  |  |  |  |  |
| PC_S1 |  |  | + | + | + | + | + | + | + |  |  |  |  |  |  |  |  |  |  |  |
|  | PC_E |  | + | + | + | + | + | + |  | + | + |  |  |  |  |  |  |  |  |  |
|  | PC_F |  | + | + | + | + | + | + |  | + | + |  |  |  |  |  |  |  |  |  |
| PC_S2 |  |  | + | + | + | + | + | + | + |  |  |  |  |  |  |  |  |  |  |  |
| PC_S3 |  |  | + | + |  |  |  |  |  |  |  |  |  |  |  |  |  |  |  |  |
| PC_S4 |  |  | + | + |  |  |  |  |  |  |  |  |  |  |  |  |  |  |  |  |
|  | PC_G |  | + | + | + | + | + | + |  | + | + |  |  |  |  |  |  |  |  |  |
|  | PC_H |  | + | + | + | + | + | + |  | + | + |  |  |  |  |  |  |  |  |  |
| PC_S5 |  |  | + | + |  |  |  |  |  |  |  |  |  |  |  |  |  |  |  |  |
| PC_S6 |  |  | + | + |  |  |  |  |  |  |  |  |  |  |  |  |  |  |  |  |
| PC_S7 |  |  | + | + |  |  |  |  |  |  |  |  |  |  |  |  |  |  |  |  |
|  | I |  | + | + | + | + |  |  | + | + | + |  |  |  |  |  |  |  |  |  |
|  | J |  | + | + | + | + |  |  | + | + | + |  |  |  |  |  |  |  |  |  |

<sup>a</sup>: Bacteriophage samples isolated by infecting *L. lactis* MG1363 with CHPC966 phage lysate. The host range of phage CHPC966 is shown in the first row of the table.
<sup>b</sup>: Bacteriophage samples isolated by infecting *L. lactis* MG1363 with a phage lysate derived from the first infection event. These isolates and the ones they were derived from are grouped and shown with a light grey background.
<sup>c</sup>: +, infected; no entry, not infected.

As shown in Table 3, not only do the 16 isolates have different host ranges, but also their host range is considerably different from that of phage CHPC966. These results support the initial idea that the phage population resulting from infection of *L. lactis* MG1363 with phage CHPC966 is composed of different bacteriophages.

### Different phage isolates have different genome sequences

To further analyze the differences between the obtained phage lysates, the genomes of a representative group of these isolates and that of phage CHPC966 were sequenced and analyzed. Lysates that showed different host ranges and morphology of the originating plaque were selected (Table 4). In three of the lysate samples, namely B1, B3, and S1, a single contig encompassing a complete c2 bacteriophage genome, was detected, while the remaining contigs represented host chromosomal DNA contamination. The phage B1 genome displayed a nucleotide sequence (nt) identity of 85% with the genome of bacteriophage CHPC966, while this percentage was 83% for the genomes of phages B3 and S1 (Figure 2, panel A; given the high percentage of sequence identity between the genomes of the phages B3 and S1, the latter was not included in the figure). Furthermore, the genomes of B3 and S1 are 99% identical at the nt level but share 93% nt identity with the phage B1 genome (data not shown). We also conducted a detailed analysis of the proteins predicted to be responsible for receptor recognition. In c2 bacteriophages these are encoded by three genes in the late transcribed region of the genome (26). A classification of c2 phages into two subgroups, named c2 and bIL67, has been proposed based on the differences in the amino acid (aa) sequences of these proteins (27). Bacteriophages of the two subspecies recognize different protein receptors on the surface of their hosts (27). Following this classification approach, we analyzed the aa sequences of the three abovementioned proteins encoded by the phages B1, B3, S1, and CHPC966 by comparing them with those of the reference phages c2 and bIL67. While phage CHPC966 was shown to belong to the bIL67 subspecies, the three new isolates appeared to be part of the c2 subspecies (data not shown). These results, together with the differences in plaque morphology and host range of the phages, proved that the newly isolated bacteriophages are different from CHPC966.

**Figure 2.**
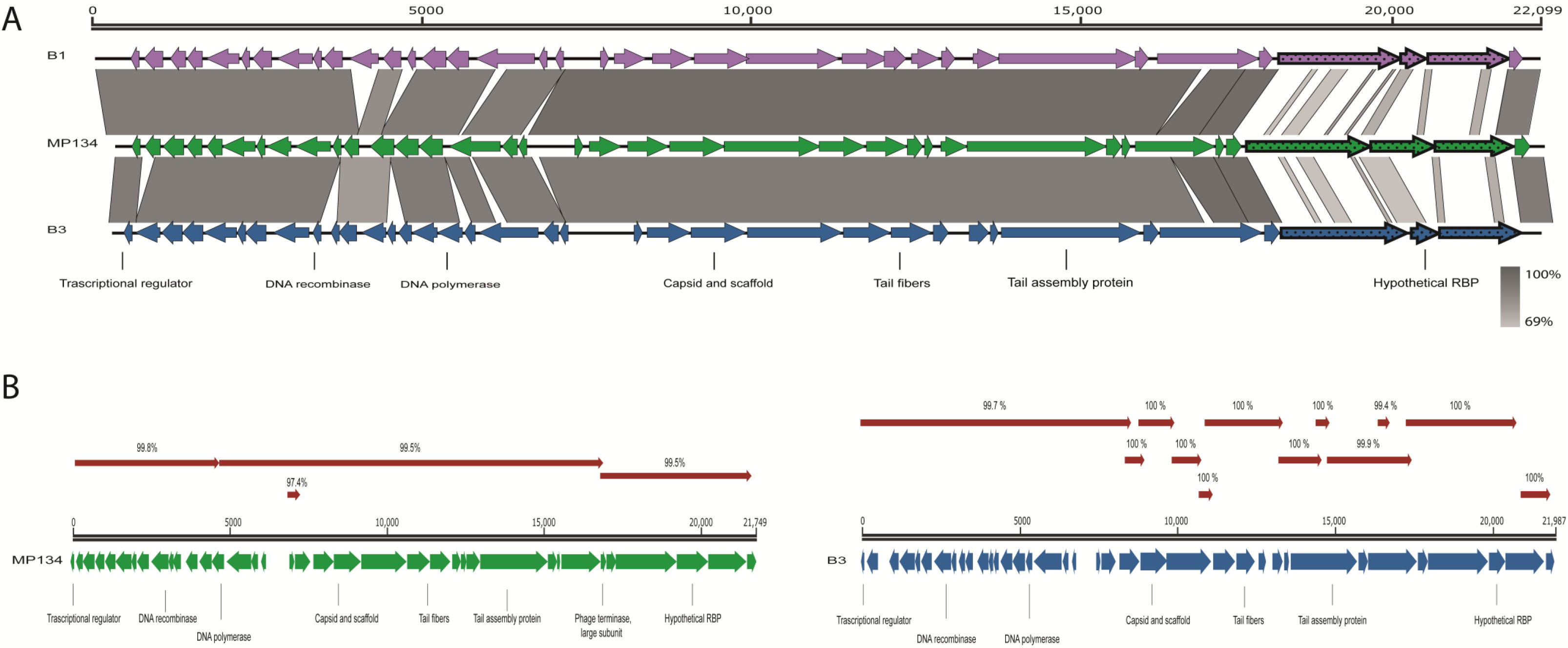
Representation of phage genomes and composition of different representative isolates studied here. Panel A: Genome comparison of phages PC_B1 and PC_B3, each separately with CHPC966. Phage genes are shown by purple, green or blue arrows. Gray boxes indicate the percentage of amino acid identity between gene products, for which a gray scale is given at the bottom right of the panel. The comparisons are to be intended as following: phage CHPC966 versus phage PC_B1, and phage CHPC966 versus phage PC_B3. Three late-transcribed genes encoding proteins predicted to be involved in host recognition are depicted with thick lines and dotted filling. Annotations for the predicted ORFs of the three phages are given at the bottom of the diagram. The genome length, in bp, is reported at the top of the diagram and refers to the phage PC_B1 genome. Panel B: Alignment between the contigs obtained from sequencing of phage PC_B9 and the genomes of phages CHPC966 and PC_B3. Phage PC_B9 different contigs are represented as red arrows. Contigs are shown only in the alignment with the phage genome with which they share the higher nt identity. The numbers above each contig indicate the percentage of nt identity between the contig and the corresponding portion of the reference genome. Nucleotide length references are given above each genome diagram, annotations for the predicted ORFs of the three phages are given at the bottom of the diagram.

**Table 4.** Bacteriophage composition of the phage lysates analyzed in this study.

| Phage name<br>(1<sup>st</sup> infection event) | Phage name<br>(2<sup>nd</sup> infection event)<sup>a</sup> | Coverage %<br>(CHPC966)<sup>b</sup> | Coverage %<br>(CSLC)<sup>c</sup> | Average % identity with<br>first-generation phage<sup>d</sup> |
| --- | --- | --- | --- | --- |
| PC_B1 |  | 0 | 100 |  |
|  | PC_A | 0 | 100 | 99 |
|  | PC_B | 0 | 100 | 99,6 |
| PC_B2 |  | 25,61 | 74,39 |  |
| PC_B3 |  | 0 | 100 |  |
| PC_B5 |  | 0,91 | 99,09 |  |
| PC_B6 |  | 0,6 | 99,4 |  |
|  | PC_C | ca. 0 | ca. 100 | 99,2 |
|  | PC_D | ca. 0 | ca. 100 | 99,5 |
| PC_B7 |  | 1,88 | 98,12 |  |
| PC_S1 |  | 0 | 100 |  |
|  | PC_E | ca. 0 | ca. 100 | 99,7 |
|  | PC_F | ca. 0 | ca. 100 | 99,5 |
| PC_S4 |  | 98,1 | 1,9 |  |
|  | PC_G | ca. 0 | ca. 100 | 99,3 |
|  | PC_H | ca. 0 | ca. 100 | 99 |
| PC_S7 |  | 99,9 | 0,01 |  |
|  | PC_I | n.a. | n.a. | 94,1 |
|  | PC_J | n.a. | n.a. | 95 |

<sup>a</sup>: Phage samples isolated during the second infection event are placed at the right of the phage samples used to generate them.
<sup>b,c</sup>: Values indicate the relative % coverage of the contigs that, in each sample, aligned better to the phage CHPC966 or the CSLC genomes, respectively, compared to the total coverage of all the contigs encoding for bacteriophage c2 genes. No values are reported for phages PC\_I and PC\_J since the small difference in nt percentage of identity of the analyzed contigs towards the genomes of phages CHPC966 and CSLC did not allow for contig classification. n.a. = not applicable.
<sup>d</sup>: Values are the average percentage of nt identity between the CSLC genome contigs of the phage isolated in the 2<sup>nd</sup> infection event and the contigs of the CSLC phage genome in the corresponding phage from the 1<sup>st</sup> infection event.

### Some of the plaques contain two different bacteriophages

Sequencing of the total DNA in the remaining phage lysates did not produce single contigs encompassing a complete c2 bacteriophage genome. Instead, many different contigs were obtained that aligned with different portions of the genome of phage CHPC966. The sequencing coverage and lengths of the contigs varied, but they were always smaller than the average length of a c2 phage genome (ca. 21 kb). Furthermore, in cases where different contigs obtained from the DNA of one lysate aligned with the same area of the phage CHPC966 genome, they would have mutually different percentages of sequence identity with that genomic region. This initial analysis suggested that the lysate samples contained more than one c2-type bacteriophage: CHPC966, and a second one, that we hypothesized to be similar to the phages in samples B1, B3 and S1.

We thus proceeded with a more detailed analysis of the various contigs contained in each sample. All contigs with a length ≥ 1000 bp and/or a coverage ≥ 1000X were selected and aligned to the genome sequences of phages CHPC966 and B3, the latter being a representative of the three already identified phage isolates. The contigs in all the samples showed different percentages of similarity with both genomes, although the vast majority of them aligned to either one or the other of the reference genomes with a higher percentage of identity. In every case, almost the entire length of the genomes of both phages CHPC966 and B3 could be covered when taking into account only those contigs that better aligned with them. Figure 2, panel B gives a representative illustration of these results. It is interesting to notice that, in several lysate samples, only the portion of the phage CHPC966 genome encoding the proteins involved in host recognition did not align with any of the sample contigs. These results underpin the initial hypothesis that the analyzed lysates contained two different c2 bacteriophages. A typical result of these alignments is reported in Table S1.

Next, the relative abundance of the two phages in each sample was determined. First, the coverage of each contig was normalized by its length, after which the total sequencing coverage of all contigs containing c2 phage genes was calculated by adding them all up. Then, the contigs were divided into two groups, based on which of the two reference genomes they showed the highest nt identity with. Finally, the total coverage of all the contigs in each group was calculated. This allowed calculating the percentage of sequencing coverage of all the contigs better aligning to either one of the phage CHPC966 or B3 genome, and, in turn, to determine which of the two phages was more represented in each sample. The results of this analysis (Table 4) show that lysates S4 and S7 were mainly composed of phage CHPC966 and that the remaining lysates (B2, and B4 through B9) mainly contained bacteriophage B3 (or a phage very similar to it). These results suggest that phage lysates deriving from plaques with a 1 mm morphology largely consist of phage CHPC966, while samples deriving from plaques with a 3 mm morphology are principally composed of a different, newly identified, c2 bacteriophage. Only sample S1 does not comply with this classification since it is derived from a 1 mm plaque but showed to be entirely composed of a phage different from CHPC966. A possible mistake in phage purification from this plaque or in labeling could be the reason for this observation.

### The newly identified bacteriophage(s) resides within the *L. lactis* MG1363 population

To identify the origin of the newly identified c2 bacteriophage(s) we first analyzed the sequencing results of phage CHPC966 to verify whether its lysate contained, even in small amounts, DNA sequences highly similar to that of the phage B3 genome. Only one contig was found that encompassed the entire genome of phage CHPC966 and all the other retrieved contigs only represented contaminating host chromosome fragments. This ruled out the possibility that the newly discovered phage was already present in the CHPC966 phage lysate used to obtain all phage isolates studied here. We then examined the only other possible source in which this phage could reside: *L. lactis* MG1363. The entire genome of the specific *L. lactis* MG1363 strain used here was sequenced and the raw nt sequences were aligned against the genome sequence of the reference phage B3. A total of 44 sequences ranging from 80 to 150 nt in length in the *L. lactis* MG1363 raw data shared nt identities between 85% and 100% with different regions of the bacteriophage B3 genome (Figure 3, Table S2). Since the lactococcal genome sequences comprised a total of 6.5 million reads, and given the genome sizes of both *L. lactis* MG1363 and bacteriophage B3 (2.5 Mb and 22 Kb, respectively) we estimated that the *L. lactis* MG1363 sample contains approximately one B3-like bacteriophage for every 270 bacterial cells. These results strongly suggest that a relationship typical of the carrier state life cycle (CSLC) exists between a c2 bacteriophage (highly similar to phage B3) and *L. lactis* MG1363. We hypothesize that infection of this strain by bacteriophage CHPC966 allows the CSLC bacteriophage to enter into an active lytic cycle and to produce visible lysis plaques on a double agar overlay plaque assay. To rule out that the newly identified phage is present in a pseudolysogenic state, the *L. lactis* MG1363 strain was colony-purified via ten consecutive single-colony propagations. The 10 purified colonies were infected with phage CHPC966 in a double agar overlay assay and in all cases plaques with the two distinct morphologies (1 and 3 mm diameter) were clearly distinguishable (data not shown). Also in these cases, the number of retrieved plaques with 1 mm morphology corresponded with the phage lysate dilution used in the assay, while the number of the 3 mm plaques remained constant (≤ 50) regardless of the lysate dilution used. These data suggest that the newly identified bacteriophage was still present in each colony. As bacterial populations can be easily cured from pseudolysogenic phages via colony purification (8, 28), our results thus strongly suggest that the newly identified bacteriophage is present in a CSLC state. Additionally, bacteriophages of the c2 species are strictly lytic as they do not possess the genetic setup to integrate into the host chromosome and cannot, therefore, establish a lysogenic or pseudolysogenic relationship with their host.

**Figure 3.**
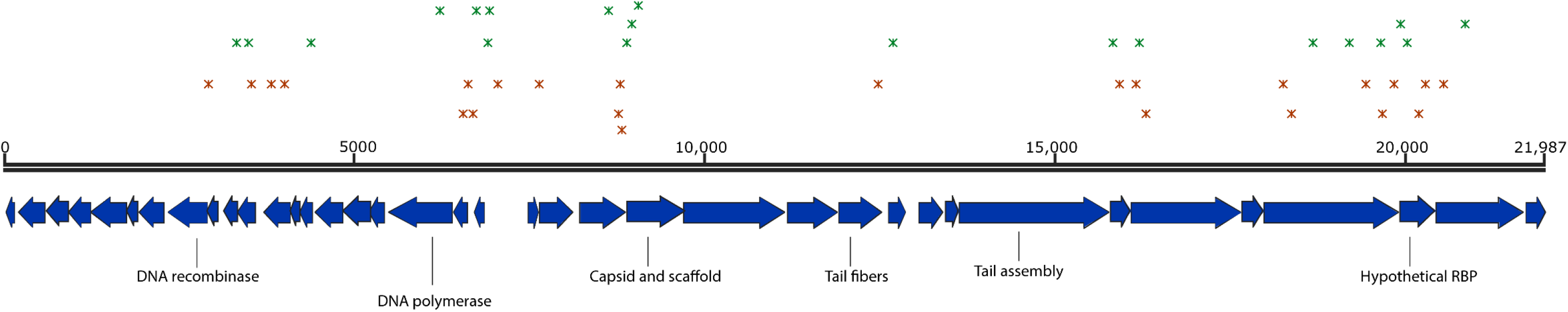
Alignment between *L. lactis* MG1363 raw genome sequencing data and the bacteriophage PC_B3 genome sequence. The bacteriophage PC_B3 genome is depicted. Annotation of genes (blue arrows) is reported at the bottom of the figure while nucleotide length reference is given above the genome. Asterisks represent the alignment starting position between the identified sequences in the raw genome sequencing data of the *L. lactis* MG1363 genome and the bacteriophage B3 genome. Green and red asterisks represent plus and minus strand alignment, respectively.

### A CSLC bacteriophage might reside in many of the *L. lactis* MG1363 samples used worldwide

To rule out that the CSLC phage is a phage contamination of the specific *L. lactis* MG1363 strain used in this work, infection with phage CHPC966 was repeated on four stocks of *L. lactis* MG1363 strain present in our laboratory and independently stored at different moments over the last decade, and on a fifth stock that we kindly received from Douwe van Sinderen laboratory, in Ireland. Additionally, the *L. lactis* MG1363 derivative NZ9000 (29) was infected with two c2 bacteriophages other than CHPC966. In all cases, plaques with the two distinct morphologies were clearly visible in each plate (Figure S1). Although the genomes of the phages obtained from these infection events were not sequenced, these results strongly suggest that a c2 bacteriophage might reside in many of the *L lactis* MG1363 derivatives used worldwide, and that infection of this strain with phages of the c2 species can induce lytic replication of the CSLC phage.

### The CSLC phage can actively propagate if employed for subsequent infections

To better understand the dynamics of phage-host interaction between the CSLC phage and *L. lactis* MG1363, phages B2, B6, and S1 (derived from plaques primarily composed of the CSLC-derived phage) and phages S4 and S7 (originating from plaques mainly composed of phage CHPC966) were used in a second round of infection of *L. lactis* MG1363. First, the morphology of the plaques originated from these infections was evaluated, and subsequently two single plaques were isolated from each infection event and propagated in liquid cultures of the same host. The new phage isolates were named CP_A through CP_J; for ease of reading we will refer to them only with the letters A through J. The host range of the phages in these newly obtained phage lysates was assessed and the phage genomic DNA was isolated and sequenced. The plaque morphology was homogeneous in all cases: all plaques are 3 mm in diameter and the number of plaques corresponded with the lysate dilution used in the assay (data not shown). Interestingly, in all cases the host range of the 10 isolates differed from that of the isolate from which they originated (Table 3). Analysis of the DNA sequence reads revealed that a single contig containing a complete c2 bacteriophage was not present in any of the samples. Subsequently, all contigs with a length ≥ 1000 bp and/or a coverage ≥ 1000X were selected and aligned to the genomes of phage CHPC966 and the CSLC phage. For phages A and B, and phages E and F, CSLC phages B1 and S1 were used, respectively, while the contigs of the other phage samples were compared with the genome of the CSLC-derived phage B3 (Table 4).

Phages A through F originated from samples mainly containing the CSLC-derived phage. Almost all their contigs showed a higher percentage of nt identity with the CSLC-derived phage genome than with that of phage CHPC966. A maximum of 3 contigs in phages C, D or F better aligned to phage CHPC966 genome. The length of these alignments was between 342 and 1384 bp and the coverage of these contigs ranged between 0.06% and 0.4% of the total coverage of contigs containing bacteriophage c2 genes. Phages G and H originated from a lysate mainly composed of phage CHPC966, although nearly all their contigs showed a higher percentage of nt identity with the phage B3 genome. In both samples, two contigs with a coverage of 0.08% to 1.4% of the total analyzed contigs coverage showed a higher percentage of identity with the genome of phage CHPC966. The total lengths of the aligned contigs ranged between 1287 and 1808 bp. In these and the previous cases, these low abundant contigs aligned with genes of phage CHPC966 that code for proteins of unknown function. Finally, phages I and J originated from a sample mainly composed of phage CHPC966. When aligned with the genome sequence of phages CHPC966 and B3, the sequenced contigs of these two samples showed higher nt identity with the latter phage.

However, the average percentage of identity (94.5) with the phage B3 genome was lower than the average percentage of identity that contigs of phages A through H showed with the CSLC phage (99.3). We hypothesize that these two samples each contain one single CSLC-derived phage, the genome of which differs from the reference CSLC genome used in the alignment, more than those of the rest of the analyzed samples. Based on these results, the infection events that generated phages G through J seem similar to the infection that generated phages B1, B3 and S1. The change in plaque morphology, namely heterogeneous in the first infection round and homogeneous in the second, might be due to the fact that in the latter cases, the CSLC-derived phage was already present, albeit at very low titration, in the lysate employed in the infection.

These results, together with those collected from the isolates generated in the first infection event, show that once the CSLC bacteriophage enters into a lytic cycle (e.g. after infection of *L. lactis* MG1363 with phage CHPC966), it is capable of actively propagating with an efficiency of plating much higher than that of the CSLC bacteriophages isolated from the first infection event of *L. lactis* MG1363 with phage CHPC966.

## Discussion

In the present work we prove that a carrier state life cycle (CSLC) between a lactococcal c2 bacteriophage and the *L. lactis* plasmid-free laboratory strain MG1363 can be established and might be present in many samples of MG1363 currently used worldwide. Our data and *in silico* analyses support the hypothesis that the CSLC phage population exists in a state of equilibrium with the *L. lactis* MG1363 population. Given the very low abundance of the CSLC phage in the *L. lactis* MG1363 population, the nt sequences corresponding to this phage genome would normally be excluded from the *L. lactis* MG1363 genome assembly process and would, therefore, be overlooked in a classic BLAST alignment between the *L. lactis* MG1363 final assembled genome sequence and the CSLC phage genome sequence. Our data further indicates that infection of *L. Lactis* MG1363 with the c2 phage CHPC966 induces the CSLC phage to actively replicate and generate visible progeny in a double agar overlay plaque assay. This infection event produces a mixed phage population composed of phage CHPC966 and the CSLC phage in different proportions. Three types of plaques are retrieved in this case. The first type, which represents the majority of the visible plaques, is mainly composed of phage CHPC966 and is 1 mm in diameter. The second type is primarily composed of the CSLC phage and can either have a diameter of 1 mm or 3 mm. Finally, there are plaques with a diameter of 3 mm that are uniquely composed of the CSLC phage. The phages composing the lysates obtained from plaques with different morphologies show considerably different host ranges and different genome sequences. Once isolated and subsequently used in a second round of infection of *L. lactis* MG1363, the CSLC-derived phage can actively replicate and infect different hosts than the phage it originated from could infect, producing plaques of homogeneous morphology (3 mm diameter). Nucleotide sequencing of the genomic DNA isolated from these plaques proved that they are almost entirely composed of the CSLC phage and that, in some cases, small portions of the phage CHPC966 genome can be found at very low concentration. The sequence of the CSLC phage isolated from these infection events appears to be different from that of the isolates they originated from. A graphic overview of these events is given in Figure 4. It has been previously postulated that CSLC bacteriophages live in intimate relationship with their host either by being externally bound to the surface of some host cells and waiting for sensitive host bacteria to become available, or by being present as an episome inside a subpopulation of cells (9, 8).

**Figure 4.**
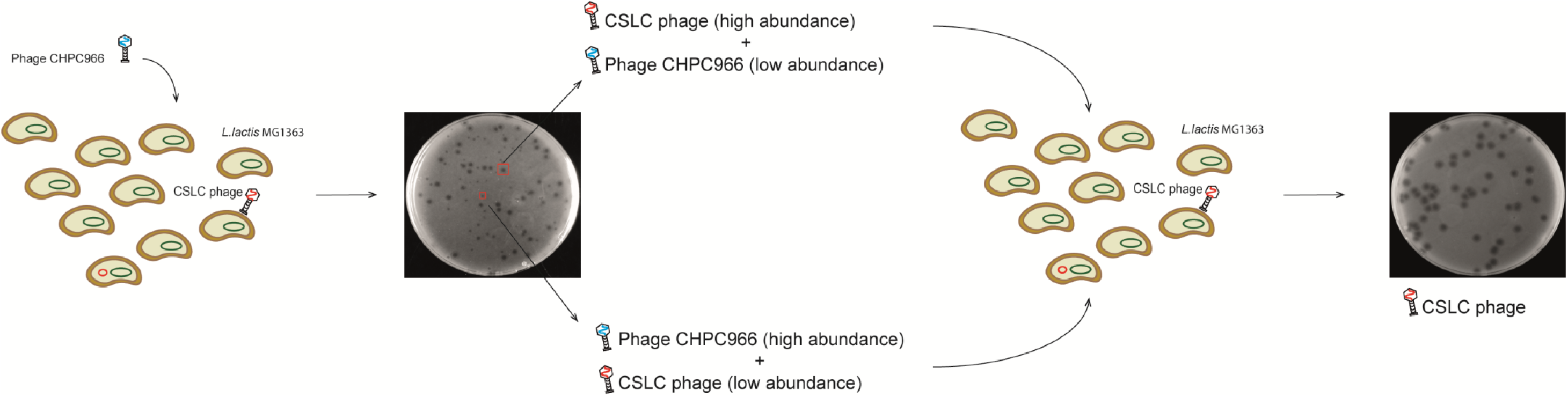
Schematic representation of the two infection events studied here and of their outcomes. One representative plate is shown for each infection event outcome and together they show the differences in plaque morphology in the two experiments. The composition of the phage population resulting from each infection event is represented with a cartoon illustration. Phages CHPC966 and CSLC are distinguished by the color of their genome: blue and red, respectively. The CSLC phage is depicted either as being bound on the surface of a subpopulation of *L. lactis* MG1363 cells, or in an episomic form (red circle). *L. lactis* MG1363 genome is depicted in green.

On the basis of this knowledge and our results, we conclude that each infection event studied here represents, in fact, a co-infection where two c2-type phages, i.e. CHPC966 and the CSLC phage, infect *L. lactis* MG1363 at different multiplicity of infection. It is known that c2 phages have very conserved genomes with specific areas that are more prone to variation (26). One of these areas contains three late-transcribed genes coding for proteins thought to be involved in host recognition (26). If, during infection of *L. lactis* MG1363, phage CHPC966 infects a cell containing the CHLC phage in an episomic form, homologous recombination could take place at specific sequences located in the highly variable portions of their genomes producing a population of CHPC966 and CSLC phages with modified receptor-binding protein-encoding genes. The modified host recognition structure, in turn, allows the recombinant CSLC phages to infect a larger portion of the lactococcal population and to release a quantifiable, albeit small, viral progeny. When the resulting CSLC phages are used to infect *L. lactis* MG1363 once more, they will be able to infect more cells in the host population and produce even more progeny. At the same time, this second co-infection event between the infecting CSLC phage and the still residing CSLC episome produces a new population of CSLC phage variants. This might represent a mechanism of hyper fast phage evolution. Although more phage isolates from a larger number of consecutive infections should be collected and analyzed to allow drawing a final conclusion on the evolutionary aspect of the events uncovered here, it is tempting to speculate that the specific c2-phage-*L. lactis* host interaction described here represents a mechanism that *L. lactis* c2 phages employ to rapidly modify their genome and to better adapt to their environment and seek out a wider number of hosts.

This is, to the best of our knowledge, the first reported study on the evolutionary implication that a CSLC relationship has on a specific bacteriophage. Our work sheds new light on what is known on phage-host interactions and infection outcomes in *L. lactis*, and on possible evolutionary strategies employed by lactococcal bacteriophages. These findings also represent an important discovery for dairy industries as they point to the importance of careful starter strains selection: the use of a starter culture containing a non-identified CSLC phage can lead to unforeseen infection bursts and to the release of recombinant phages in the production vats. Finally, we believe that this work provides pioneering results and valuable insights into alternative phage-host interactions in Gram-positive bacteria and demonstrates that phage infection outcomes can be more intricate than currently postulated.

## Aknowledgements

We thank Danny Incarnato for valuable discussions. This work was carried out within the BE-Basic R&D Program, which was granted an FES subsidy from the Dutch Ministry of Economic Affairs.

