## Supplemental material Table 1 for "Alternative phage-host interaction in *Lactococcus lactis*: the carrier state drives rapid evolution of phages"

**Table S1: Representative results of the alignment of the contigs obtained from sequencing of isolate PH_B9 with the genome of the CSLC phage and of phage CHPC966.**

| contig number[a] | Subject[b] | % nt identity | alignment lenght (bp) | mismatches | gaps | Subject_start[c] | Subject_end[d] | Evalue |
|---|---|---|---|---|---|---|---|---|
| 1 | CHPC966 | 99,282 | 9605 | 65 | 1 | 6451 | 16051 | 0 |
| 1 | CHPC966 | 100 | 1010 | 0 | 0 | 5429 | 6438 | 0 |
| 1 | CHPC966 | 100 | 654 | 0 | 0 | 16052 | 16705 | 0 |
| 1 | CHPC966 | 99,78 | 455 | 1 | 0 | 4932 | 5386 | 0 |
| 1 | CHPC966 | 98,678 | 454 | 6 | 0 | 16706 | 17159 | 0 |
| 3 | CHPC966 | 99,502 | 4816 | 19 | 4 | 21728 | 16916 | 0 |
| 4 | CHPC966 | 99,893 | 4672 | 5 | 0 | 1 | 4672 | 0 |
| 79 | CHPC966 | 97,494 | 399 | 10 | 0 | 7310 | 6912 | 0 |
| 2 | PC_B3 | 99,981 | 5234 | 1 | 0 | 11 | 5244 | 0 |
| 2 | PC_B3 | 99,285 | 2519 | 0 | 1 | 5306 | 7824 | 0 |
| 2 | PC_B3 | 100 | 824 | 0 | 0 | 7868 | 8691 | 0 |
| 2 | PC_B3 | 100 | 1034 | 0 | 0 | 20954 | 21987 | 0 |
| 5 | PC_B3 | 100 | 3540 | 0 | 0 | 17562 | 21101 | 0 |
| 6 | PC_B3 | 100 | 2170 | 0 | 0 | 13297 | 11128 | 0 |
| 6 | PC_B3 | 100 | 328 | 0 | 0 | 13622 | 13295 | 1,09E-175 |
| 12 | PC_B3 | 100 | 1371 | 0 | 0 | 13475 | 14845 | 0 |
| 13 | PC_B3 | 99,91 | 1110 | 0 | 1 | 17709 | 16601 | 0 |
| 13 | PC_B3 | 100 | 1076 | 0 | 0 | 16469 | 15394 | 0 |
| 13 | PC_B3 | 100 | 281 | 0 | 0 | 15283 | 15003 | 1,53E-149 |
| 16 | PC_B3 | 100 | 1154 | 0 | 0 | 9017 | 10170 | 0 |
| 20 | PC_B3 | 100 | 958 | 0 | 0 | 10023 | 10980 | 0 |
| 22 | PC_B3 | 87,528 | 890 | 106 | 4 | 5244 | 4359 | 0 |
| 38 | PC_B3 | 100 | 621 | 0 | 0 | 9164 | 8544 | 0 |
| 65 | PC_B3 | 100 | 453 | 0 | 0 | 14698 | 15150 | 0 |
| 67 | PC_B3 | 100 | 443 | 0 | 0 | 11275 | 10833 | 0 |
| 88 | PC_B3 | 99,475 | 381 | 2 | 0 | 11109 | 10729 | 0 |
| 94 | PC_B3 | 99,469 | 377 | 2 | 0 | 16982 | 16606 | 0 |

| | | | | | | | | |
|---|---|---|---|---|---|---|---|---|
| **109** | **PC_B3** | **99,437** | **355** | **2** | **0** | **10751** | **11105** | **0** |
| **111** | **PC_B3** | **99,43** | **351** | **2** | **0** | **14898** | **15248** | **0** |
| **113** | **PC_B3** | **100** | **49** | **0** | **0** | **15592** | **15544** | **1,94E-21** |

[a] : Number of the contigs obtained from sequencing of the bacteriophage indicated at the top of each table section

[b] : Name of the bacteriophage the genome that was aligned with the indicated contigs

[c,d] : Nucleotide position, in the subject genome, for the start and end of the alignment, respectively.
