## Supplemental material Table 2 for "Alternative phage-host interaction in *Lactococcus lactis*: the carrier state drives rapid evolution of phages"

**Table S2. Results of the alignment between *L.lactis* MG1363 raw genome sequencing data, and bacteriophage PC_PC_B3 genome sequence.**

| Sequence number[a] | Alignment starting position[b] | strand | Alignment length (bp) | % nt identity | Numbers of mismatches | Numbers of gaps | e-value |
|---|---|---|---|---|---|---|---|
| 1 | 2922 | + | 151 | 99.338 | 1 | 0 | 4.15E-82 |
| 2 | 3322 | - | 106 | 94.34 | 6 | 0 | 2.55E-43 |
| 3 | 3491 | - | 60 | 83.333 | 10 | 0 | 2.54E-06 |
| 4 | 3527 | + | 151 | 100 | 0 | 0 | 1.70E-84 |
| 5 | 3804 | + | 128 | 87.5 | 16 | 0 | 1.41E-32 |
| 6 | 4006 | + | 151 | 96.689 | 5 | 0 | 1.46E-72 |
| 7 | 4374 | - | 74 | 87.838 | 9 | 0 | 4.61E-17 |
| 8 | 6221 | - | 34 | 88.235 | 4 | 0 | 3.96E-05 |
| 9 | 6542 | + | 151 | 100 | 0 | 0 | 1.70E-84 |
| 10 | 6610 | + | 151 | 99.338 | 1 | 0 | 4.15E-82 |
| 11 | 6691 | + | 150 | 99.333 | 1 | 0 | 1.64E-81 |
| 12 | 6749 | - | 51 | 92.157 | 4 | 0 | 2.84E-15 |
| 13 | 6914 | - | 150 | 85.333 | 22 | 0 | 2.19E-31 |
| 14 | 6930 | - | 108 | 84.259 | 17 | 0 | 2.95E-18 |
| 15 | 7049 | + | 131 | 91.603 | 9 | 1 | 1.63E-44 |
| 16 | 7628 | + | 147 | 91.837 | 12 | 0 | 1.83E-53 |
| 17 | 8622 | - | 151 | 84.768 | 23 | 0 | 1.35E-29 |
| 18 | 8756 | + | 131 | 94.656 | 7 | 0 | 7.49E-56 |
| 19 | 8780 | + | 151 | 84.768 | 23 | 0 | 1.35E-29 |
| 20 | 8807 | + | 151 | 99.338 | 1 | 0 | 4.15E-82 |
| 21 | 8886 | - | 87 | 94.253 | 5 | 0 | 2.28E-34 |
| 22 | 8949 | - | 28 | 96.429 | 1 | 0 | 1.04E-08 |
| 23 | 9044 | - | 147 | 95.918 | 6 | 0 | 8.70E-68 |
| 24 | 12484 | + | 132 | 90.909 | 10 | 2 | 9.70E-40 |
| 25 | 12687 | - | 112 | 86.607 | 15 | 0 | 2.04E-25 |
| 26 | 15834 | - | 125 | 90.4 | 12 | 0 | 2.46E-40 |
| 27 | 15914 | + | 151 | 96.026 | 6 | 0 | 3.57E-70 |
| 28 | 16162 | + | 151 | 89.404 | 16 | 0 | 2.65E-46 |

| | | | | | | | |
|---|---|---|---|---|---|---|---|
| 29 | 16202 | - | 151 | 96.689 | 5 | 0 | 1.46E-72 |
| 30 | 16295 | + | 151 | 99.338 | 1 | 0 | 4.15E-82 |
| 31 | 18263 | + | 151 | 92.715 | 11 | 0 | 3.07E-58 |
| 32 | 18371 | + | 151 | 100 | 0 | 0 | 1.70E-84 |
| 33 | 18671 | - | 88 | 86.364 | 12 | 0 | 2.95E-18 |
| 34 | 19198 | - | 151 | 88.079 | 18 | 0 | 1.57E-41 |
| 35 | 19439 | + | 145 | 93.103 | 10 | 0 | 4.80E-57 |
| 36 | 19646 | - | 122 | 88.525 | 14 | 0 | 9.00E-34 |
| 37 | 19678 | + | 151 | 99.338 | 1 | 0 | 4.15E-82 |
| 38 | 19834 | + | 136 | 91.912 | 8 | 1 | 4.29E-48 |
| 39 | 19937 | - | 52 | 86.538 | 7 | 0 | 1.04E-08 |
| 40 | 20031 | - | 127 | 82.677 | 22 | 0 | 1.17E-17 |
| 41 | 20193 | + | 150 | 99.333 | 1 | 0 | 1.64E-81 |
| 42 | 20286 | + | 151 | 91.391 | 13 | 0 | 1.83E-53 |
| 43 | 20548 | + | 128 | 90.625 | 12 | 0 | 3.98E-42 |
| 44 | 20850 | - | 130 | 86.923 | 17 | 0 | 2.19E-31 |

[a] : Sequences identified in the *L.lactis* MG1363 raw genome sequencing data that align with bacteriophage PC_PC_B3 genome sequence.

[b] : Nucleotide position in bacteriophage PC_PC_B3 genome corresponding to the alignment start.
