## Supplemental material Figure S1 for "Alternative phage-host interaction in *Lactococcus lactis*: the carrier state drives rapid evolution of phages"

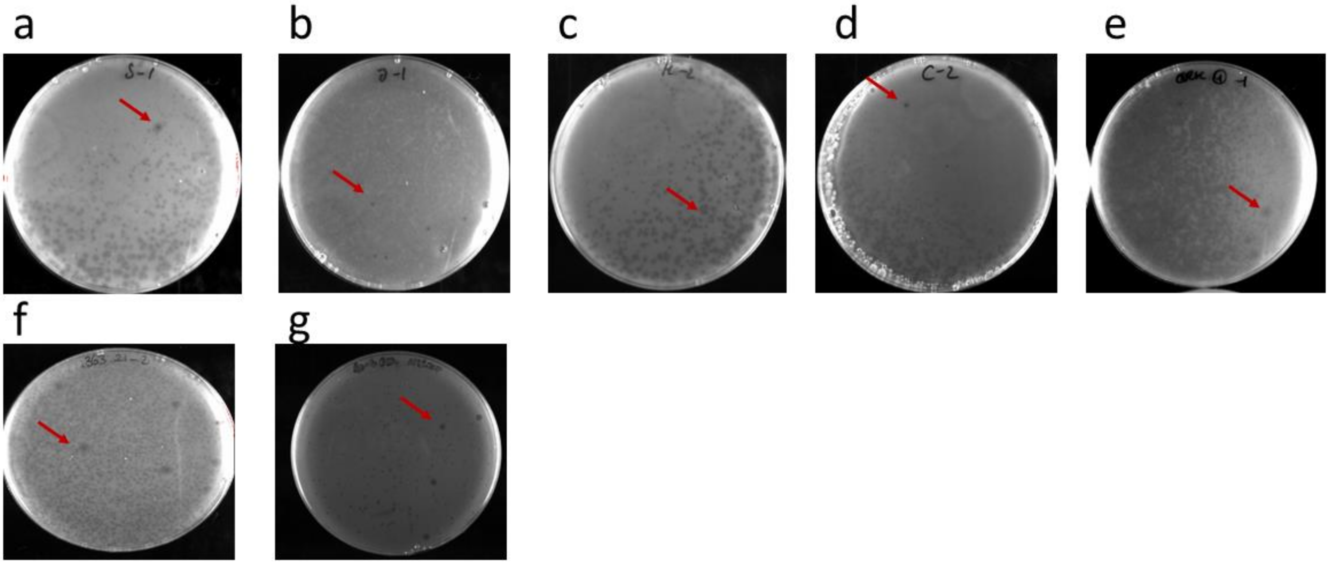


**Figure S1. comparison of plaque morphology from different infections of *L. lactis* MG1363 and its derivative NZ9000.**

Panels a through e: plaques obtained by infecting four different *L. lactis* MG1363 strain stocks stored in our laboratory in different moments over the last decade, with phage CHPC966. Panel d: plaques obtained by infecting a *L. lactis* MG1363 stock strain from a laboratory in Ireland, with phage CHPC966. Panels e and f: plaques obtained by infecting *L. lactis* strains NZ900 with two c2 bacteriophages different from each other and from phage CHPC966. The red arrows indicate the plaques with a 3mm diameter morphology that are presumably composed of the CSLC phage.
